# Genome-wide association study of suicide attempt in psychiatric disorders identifies association with major depression polygenic risk scores

**DOI:** 10.1101/416008

**Authors:** Niamh Mullins, Tim B. Bigdeli, Anders D Børglum, Jonathan R I Coleman, Ditte Demontis, Ayman H. Fanous, Divya Mehta, Robert A. Power, Stephan Ripke, Eli A Stahl, Anna Starnawska, Adebayo Anjorin, Aiden Corvin, Alan R Sanders, Andreas J Forstner, Andreas Reif, Anna C Koller, Beata Świątkowska, Bernhard T Baune, Bertram Müller-Myhsok, Bettina Konte, Brenda WJH Penninx, Carlos Pato, Clement Zai, Dan Rujescu, Digby Quested, Douglas F Levinson, Elisabeth B Binder, Enda M Byrne, Esben Agerbo, Fabian Streit, Fermin Mayoral, Frank Bellivier, Franziska Degenhardt, Gerome Breen, Gunnar Morken, Gustavo Turecki, Guy A Rouleau, Hans J Grabe, Henry Völzke, Ina Giegling, Ingrid Agartz, Ingrid Melle, Jacob Lawrence, James B Potash, James TR Walters, Jana Strohmaier, Jianxin Shi, Joanna Hauser, Joanna M Biernacka, John B Vincent, John Kelsoe, John S Strauss, Jolanta Lissowska, Jonathan Pimm, Jordan W Smoller, José Guzman Parra, Klaus Berger, Laura J Scott, M. Helena Azevedo, Maciej Trzaskowski, Manolis Kogevinas, Marcella Rietschel, Marco Boks, Marcus Ising, Maria Grigoroiu-Serbanescu, Marian L Hamshere, Marion Leboyer, Mark Frye, Markus M Nöthen, Martin Alda, Martin Preisig, Merete Nordentoft, Michael Boehnke, Michael C O’Donovan, Michael J Owen, Michele T Pato, Miguel Renteria, Monika Budde, Myrna M Weissman, Naomi R Wray, Nicholas Bass, Olav B Smeland, Ole A Andreassen, Ole Mors, Pablo V Gejman, Pamela Sklar, Patrick McGrath, Per Hoffmann, Peter McGuffin, Phil H Lee, René S Kahn, Roel A Ophoff, Rolf Adolfsson, Sandra Van der Auwera, Srdjan Djurovic, Stanley I Shyn, Stefan Kloiber, Stefanie Heilmann-Heimbach, Stéphane Jamain, Steven P Hamilton, Susan L McElroy, Susanne Lucae, Sven Cichon, Thomas G Schulze, Thomas Hansen, Thomas Werge, Tracy M Air, Vishwajit Nimgaonkar, Vivek Appadurai, Wiepke Cahn, Yuri Milaneschi, Major Depressive Disorder Working Group of the Psychiatric Genomics Consortium, Bipolar Disorder Working Group of the Psychiatric Genomics Consortium, Schizophrenia Working Group of the Psychiatric Genomics Consortium, Kenneth S Kendler, Andrew McQuillin, Cathryn M Lewis

## Abstract

**Objective:** Over 90% of suicide attempters have a psychiatric diagnosis, however twin and family studies suggest that the genetic etiology of suicide attempt (SA) is partially distinct from that of the psychiatric disorders themselves. Here, we present the largest genome-wide association study (GWAS) on suicide attempt using major depressive disorder (MDD), bipolar disorder (BIP) and schizophrenia (SCZ) cohorts from the Psychiatric Genomics Consortium.

**Method:** Samples comprise 1622 suicide attempters and 8786 non-attempters with MDD, 3264 attempters and 5500 non-attempters with BIP and 1683 attempters and 2946 non-attempters with SCZ. SA GWAS were performed comparing attempters to non-attempters in each disorder followed by meta-analysis across disorders. Polygenic risk scoring investigated the genetic relationship between SA and the psychiatric disorders.

**Results:** Three genome-wide significant loci for SA were found: one associated with SA in MDD, one in BIP, and one in the meta-analysis of SA in mood disorders. These associations were not replicated in independent mood disorder cohorts from the UK Biobank and *i*PSYCH. Polygenic risk scores for major depression were significantly associated with SA in MDD (P=0.0002), BIP (P=0.0006) and SCZ (P=0.0006).

**Conclusions:** This study provides new information on genetic associations and the genetic etiology of SA across psychiatric disorders. The finding that polygenic risk scores for major depression predict suicide attempt across disorders provides a possible starting point for predictive modelling and preventative strategies. Further collaborative efforts to increase sample size hold potential to robustly identify genetic associations and gain biological insights into the etiology of suicide attempt.

## Introduction

Suicide is a worldwide public health problem with over 800,000 deaths due to suicide each year (1). It is the second leading cause of death among young adults and rates of suicide are far exceeded by suicide attempts, which occur up to 20 times more frequently (1). This represents a huge personal, social and economic burden, with the Centers for Disease Control and Prevention reporting that suicide costs the US economy $51 billion per year in healthcare and work-loss related costs (2). These stark figures highlight the urgent need for improved prevention and treatment, however progress has been hampered by the lack of reliable methods for predicting suicidality and a poor understanding of its biological etiology.

Over 90% of suicide attempters or victims have a psychiatric disorder, particularly mood disorders, schizophrenia and substance use disorders (3, 4). Heritability estimates of suicidal behavior from twin studies range from 30-55% and twin and family studies suggest that the genetic etiology of suicide attempt is partially distinct from that of the psychiatric disorders themselves (5, 6). Several genome-wide association studies (GWAS) have been conducted on suicide attempt, by comparing attempters versus non-attempters with depression or bipolar disorder, to test for genetic variants contributing independently to suicide attempt (7-10). These studies have failed to identify any replicable genetic associations, likely due to limited sample sizes that were underpowered to detect the genetic effects typical for a single SNP. Other GWAS have examined subjects recruited specifically on the basis of suicide attempt, or suicide attempters and non-attempters from population-based cohorts, but to date no loci have been robustly implicated (11-13).

Genetic studies have indicated that suicide attempt has a polygenic architecture, as polygenic risk scores for suicide attempt have shown modest predictive ability in independent samples and small but significant SNP-heritability estimates for suicide attempt have been reported (10, 13). These findings are consistent with the presence of small genetic effects that the original GWAS were underpowered to detect at genome-wide significance. In the current study, we present the largest GWAS on suicide attempt to date, comparing a total of 6,569 suicide attempters and 17,232 non-attempters, with major depressive disorder (MDD), bipolar disorder (BIP) or schizophrenia (SCZ) from the Psychiatric Genomics Consortium.

## Method

### Subjects and phenotype definition

Subjects were drawn from 16 MDD cohorts, 21 BIP cohorts and 9 SCZ cohorts in the Psychiatric Genomics Consortium (PGC), where information on suicide attempt (SA) had been collected. Only cases affected with psychiatric disorders were included, and all psychiatric disorders were defined using structured psychiatric interviews according to international consensus criteria (DSM-IV, ICD-9, or ICD-10) (14-16). Supplementary Tables 1-3 summarise the source, inclusion and exclusion criteria in each cohort. All subjects were of European ancestry. Suicide attempt (SA) was defined in each cohort using items from structured clinical interviews (Supplementary Table 4). Lifetime suicide attempt was defined across cohorts as a deliberate act of self-harm with at least some intent to result in death. Individuals missing information on suicide attempt were excluded. Across the MDD, BIP and SCZ datasets, there were a total of 6,569 suicide attempters and 17,232 non-attempters (Table 1). All subjects gave written informed consent to participate in the source studies.

**Table 1:**
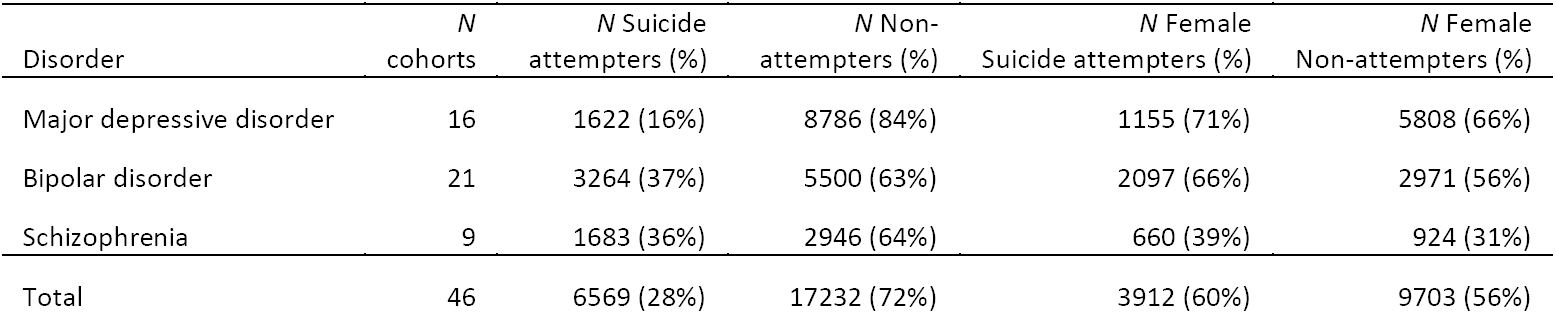
Sample characteristics from PGC cohorts

### Genotyping, quality control and imputation

Cohorts were genotyped following their local protocols, after which standardised quality control and imputation were performed centrally using the PGC ‘Ricopili’ pipeline (https://sites.google.com/a/broadinstitute.org/ricopili/), for each cohort separately. These procedures have been described in detail previously (17-19). Briefly, the quality control parameters for retaining SNPs and subjects were: SNP missingness < 0.05 (before sample removal), subject missingness < 0.02, autosomal heterozygosity deviation (F_het_ < 0.2), SNP missingness < 0.02 (after sample removal), difference in SNP missingness between psychiatric cases and healthy controls < 0.02, SNP Hardy-Weinberg equilibrium (*P* > 10^-10^ in psychiatric cases) and minor allele frequency ≥ 0.01. Genotype imputation was performed using the pre-phasing/ imputation stepwise approach implemented in IMPUTE2/ SHAPEIT (chunk size of 3 Mb and default parameters) to the 1000 Genomes Project reference panel (20-22). SNPs with an imputation INFO-score < 0.6 were excluded. The numbers of SNPs analysed were 8482392, 8807006 and 8814543 in the MDD, BIP, and SCZ datasets respectively.

### Statistical analysis

GWAS on suicide attempt were performed using PLINK 1.9 by comparing imputed marker dosages under an additive logistic regression model between suicide attempters and non-attempters in each cohort separately (23). The first five principal components (PCs), generated using EIGENSTRAT were used as covariates in all GWAS to control for population stratification (24). There was no evidence of stratification artifacts or uncontrolled test statistic inflation in the results from any cohort (e.g. λ_GC_ was 0.87 - 1.01). Within each psychiatric disorder, meta-analysis was performed using an inverse variance-weighted fixed effects model in METAL, to obtain GWAS results for suicide attempt in MDD, suicide attempt in BIP and suicide attempt in SCZ (25). A fixed effects meta-analysis was also conducted for a GWAS of suicide attempt in all three psychiatric disorders, in mood disorders only, and for BIP and SCZ.

Polygenic risk scoring was used to investigate the genetic relationship between suicide attempt and the psychiatric disorders and to test for overlap in the genetic etiology of suicide attempt in MDD, BIP and SCZ. PRSice software was used to generate polygenic risk scores (PRS), according to standard protocol (26). The GWAS results from each discovery study were pruned for linkage disequilibrium (LD) using the P value informed clumping method in PLINK (--clump-p1 1 --clump-p2 1 --clump-r2 0.1 --clump-kb 250). This preferentially retains SNPs with the strongest evidence of association and removes SNPs in LD (r^2^ > 0.1) that show weaker evidence of association within 250Kb windows, based on the LD structure in the test dataset. Subsets of SNPs were selected from the results at nine increasingly liberal P value thresholds (P < 0.0001, P < 0.001, P < 0.01, P < 0.05, P < 0.1, P < 0.2, P < 0.3, P < 0.4, P < 0.5). In the test datasets, the SNP probabilities were converted to best-guess data with a genotype call probability cut-off of 0.8. Sets of alleles, weighted by their log odds ratios (OR) from the discovery GWAS, were summed into PRS for each individual in the test datasets using PLINK. The regression model also included five PCs and a covariate for each cohort in the test dataset. The amount of variance explained by the PRS was calculated as Nagelkerke’s pseudo-R^2^.

First, PRS for BIP, major depression and SCZ were used to investigate whether genetic liability for psychiatric disorders differs between suicide attempters and non-attempters. To ensure no overlap between the discovery and test datasets, PRS were generated using PGC cohorts not included in the suicide attempt analyses. All cohorts have been described in previous GWAS on the psychiatric disorders conducted by the PGC (17-19). For major depression, meta-analyses of PGC cohorts, deCODE, GERA, *i*PSYCH, Generation Scotland and UK Biobank were available, excluding each of the 16 SA cohorts in turn. The phenotype analysed includes clinically defined MDD cases as well as self-reported MDD symptoms or treatment and is referred to as ‘major depression’ (17). These discovery GWAS had approximately 59,000 cases and 112,000 controls. The discovery GWAS for BIP consisted of 11 PGC cohorts totaling 8,711 BIP cases and 15,283 controls, and for SCZ included 25,756 SCZ cases and 35,686 controls from 40 PGC cohorts. The PRS for each psychiatric disorder were tested for association with suicide attempter versus non-attempter status in the same disorder, using the method previously described. Based on the results of these analyses, PRS for major depression were also tested for association with suicide attempt in BIP and SCZ.

Second, the results of the three GWAS on suicide attempt (SA in MDD, SA in BIP and SA in SCZ) were used in turn as discovery studies and the remaining two disorders were used as separate test datasets, to investigate whether PRS for suicide attempt in one disorder are associated with suicide attempt in another. In total, six independent hypotheses were tested using polygenic risk scoring and the Bonferroni corrected significance threshold is 0.008.

The variance in suicide attempt explained by common SNPs (SNP-heritability, 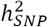) was assessed using genomic-relatedness-based restricted maximum-likelihood (GREML), implemented in GCTA software (27). The SNP probabilities were converted to best-guess data with a genotype call probability cut-off of 0.8. HapMap 3 SNPs with an INFO score ≥ 0.6 were used to calculate the genetic relatedness matrix (GRM) using PLINK 1.9, including individuals with relatedness < 0.05 (23). Ancestry informative principal components were calculated using GCTA (27). The GRM was based on a total of 1166347 SNPs in the MDD dataset, 1172705 SNPs in the BIP dataset and 1143070 SNPs in the SCZ dataset. Covariates included 20 PCs calculated using GCTA (because GRM-based analyses are more sensitive to population stratification than polygenic scoring analyses) and a covariate for each cohort within a disorder. The 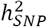 of SA in each psychiatric disorder was calculated using GCTA (27).

### Power calculations

The Genetic Power Calculator was used to determine the power to detect associations at genome-wide significance (P < 5 x 10^-8^), for the meta-analysis of 6,569 suicide attempters and 17,232 non-attempters across the three psychiatric disorders (28). This analysis had 78% power to detect an allele with frequency 0.2 and effect size 1.1 at genome-wide significance. From the GCTA-GREML power calculator, the power to detect a 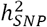 of 20% (approximately half the twin heritability estimate) for suicide attempt was 81%, 92% and 43% in the MDD, BIP and SCZ datasets respectively (29). The statistical power of polygenic risk scoring was calculated using AVENGEME software (30, 31). The power of the PRS for the psychiatric disorders to predict SA in same disorder was 51% in BIP, 93% in MDD and 78% in SCZ, given the 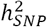for each psychiatric disorder calculated from the summary statistics (20%, 10% and 25% respectively) and hypothesizing a genetic correlation of 0.5 between the psychiatric disorder and suicide attempt. Given a 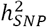of 20% for suicide attempt, the power of polygenic risk scores for suicide attempt to detect a significant difference between attempters and non-attempters in the test datasets ranged from 32-64%. The significance threshold for all polygenic risk scoring power calculations was 0.008.

### Replication

Genome-wide significant associations with suicide attempt were tested for replication in two independent mood disorder cohorts from the UK Biobank and *i*PSYCH. The UK Biobank is a population-based prospective study of 501,726 individuals, recruited at 23 centres across the United Kingdom (32). Extensive phenotypic data are available for UK Biobank participants from health records and questionnaires, including an online follow-up questionnaire focusing on mental health (Mental Health Questionnaire, MHQ [Resource 22 on http://biobank.ctsu.ox.ac.uk]). Participants were classified as having a mood disorder if they either self-reported a professional diagnosis of depression or bipolar disorder as part of the MHQ [UK Biobank field 20544, responses 10 or 11] or if they met criteria for depression or bipolar disorder on MHQ questions derived from the Composite International Diagnostic Interview (CIDI). Suicide attempters with mood disorders (n=2,149, 91% with depression) were defined as those who answered yes to the question “Have you ever harmed yourself with the intention to end your life?” [UK Biobank field f20483]. Non-attempters with mood disorders were defined as those who reported no self-harm on the MHQ (n=35,912). Genetic associations with suicide attempt were tested by comparing suicide attempters versus non-attempters with mood disorders using BGenie v.1.2 (33), covarying for six PCs and factors capturing site of recruitment and genotyping batch. Full details of the genetic quality control, imputation and mood disorder criteria are available in the Supplementary Materials.

In the *i*PSYCH cohort, individuals with mood disorders were identified based on ICD-10 codes (F30-F39) from the Danish Psychiatric Central Research Register and the National Registry of Patients, both complete until December 31, 2016 (34). Suicide attempters with mood disorders (n=4,943, 94% with MDD) were defined as those with diagnoses of suicide attempt (ICD-10: X60-X84, equivalent to intentional self-harm), those with suicide attempt indicated as ‘reason for contact’, and with a main diagnosis of poisoning (ICD-10: T39, T42, T43, and T58) or those with a diagnosis in the ICD-10: F chapter as main diagnosis and report of poisoning by drugs or other substances (ICD-10: T36–T50, T52–T60) or injuries to hand, wrist, and forearm (ICD-10: S51, S55, S59, S61, S65, S69). Individuals who died by suicide according to the Cause of Death Register were also included in the suicide attempter group. Non-attempters were defined as mood disorder cases not fulfilling any of these criteria (n=15,849). Genetic associations with suicide attempt were tested by comparing suicide attempters versus non-attempters with mood disorders, including 10 PCs and genotyping batch as covariates. Full details of quality control in the *i*PSYCH cohort are included in the Supplementary Materials.

### Results

### Sample characteristics

The proportion of psychiatric disorder cases reporting suicide attempt ranged from 16% in MDD to 36-37% in bipolar disorder and schizophrenia (Table 1). For each disorder, there was a higher proportion of females in the suicide attempters than the non-attempters. Table 1 shows the number and proportion of suicide attempters and non-attempters within each psychiatric disorder. The numbers in the individual cohorts are shown in Supplementary Tables 5-7.

### Genome-wide association studies

A GWAS of suicide attempters versus non-attempters was performed in the MDD, BIP and SCZ datasets separately. In the analysis of suicide attempt in MDD, one SNP reached genome-wide significance: rs45593736, P = 2.61 x 10^-8^, OR A allele = 2.38 (Table 2). This SNP is in an intron of the *ARL5B* (ADP-Ribosylation Factor-Like 5B) gene, and the A allele has a frequency of 0.02. In the GWAS of suicide attempt in BIP, an insertion-deletion polymorphism on chromosome 4 met genome-wide significance: chr4_23273116_D, P = 1.15×10^-8^, OR for the deletion = 1.29 (Table 2). This is an intronic variant in the non-coding RNA *LOC105374524*. In the analysis of suicide attempt in SCZ, there were no SNPs reaching genome-wide significance, but this analysis had the smallest total sample size.

**Table 2:**
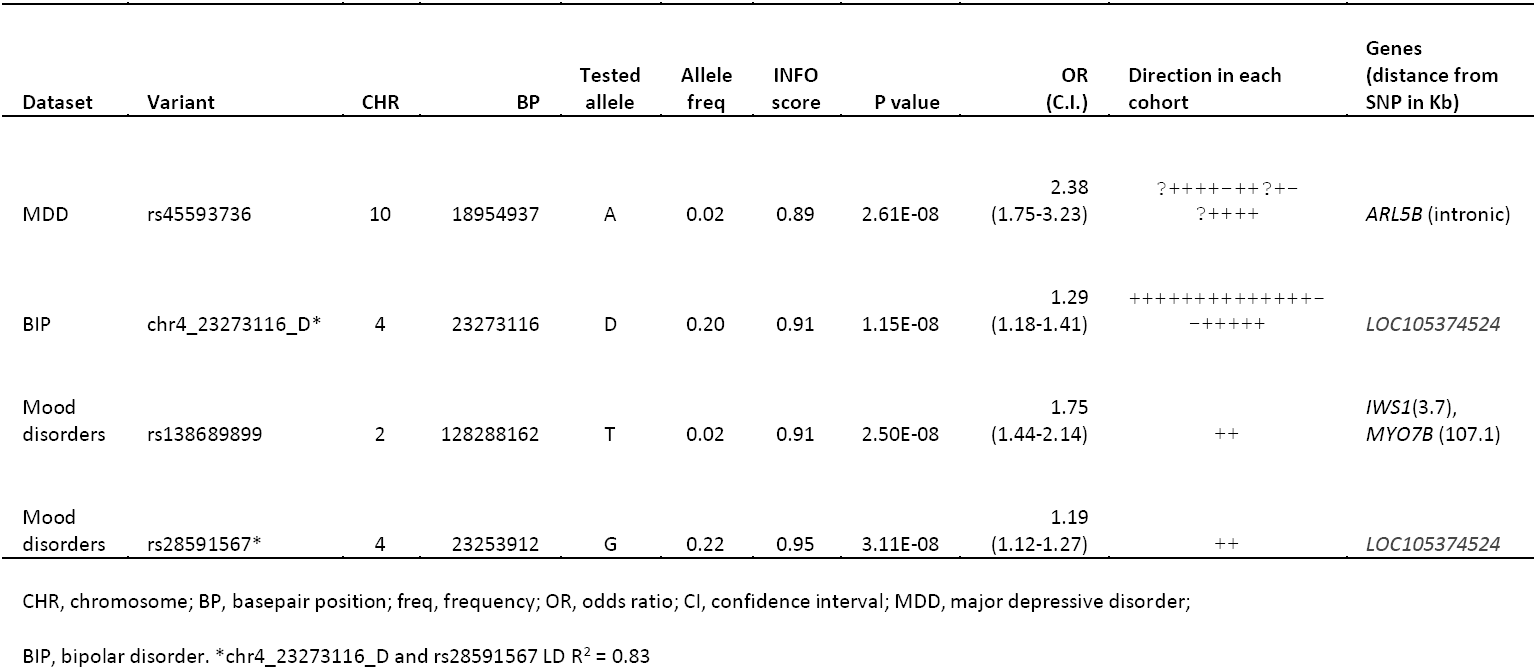
Genome-wide significant loci for suicide attempt showing the most significant variant from each genomic region

Meta-analysis of the GWAS results for suicide attempt across all three disorders produced no genome-wide significant results (Supplementary Table 11). In a meta-analysis of suicide attempt in mood disorders (MDD and BIP), there were 10 genome-wide significant SNPs from two independent genomic regions (Table 2, Figure 1). The most significant association was rs138689899 on chromosome 2, P = 2.50 x 10^-8^, OR T allele = 1.75. This is an intergenic SNP that lies between the *IWS1* and *MYO7B* genes. The other significant locus was on chromosome 4 in *LOC105374524*, as found in the BIP SA analysis. The most significant SNP was rs28591567 (P = 3.11 x 10^-8^, OR G allele = 1.19, frequency G allele = 0.22), in high LD (R^2^= 0.83) with the insertion-deletion polymorphism identified in SA in BIP (Table 2). Weak evidence for association was found in MDD (rs28591567, P = 0.03, OR allele G allele = 1.11); the locus was not associated with SA in schizophrenia (P = 0.67). No significant associations were identified in the meta-analysis between SA in BIP and SCZ (Supplementary Table 13).

**Figure 1:**
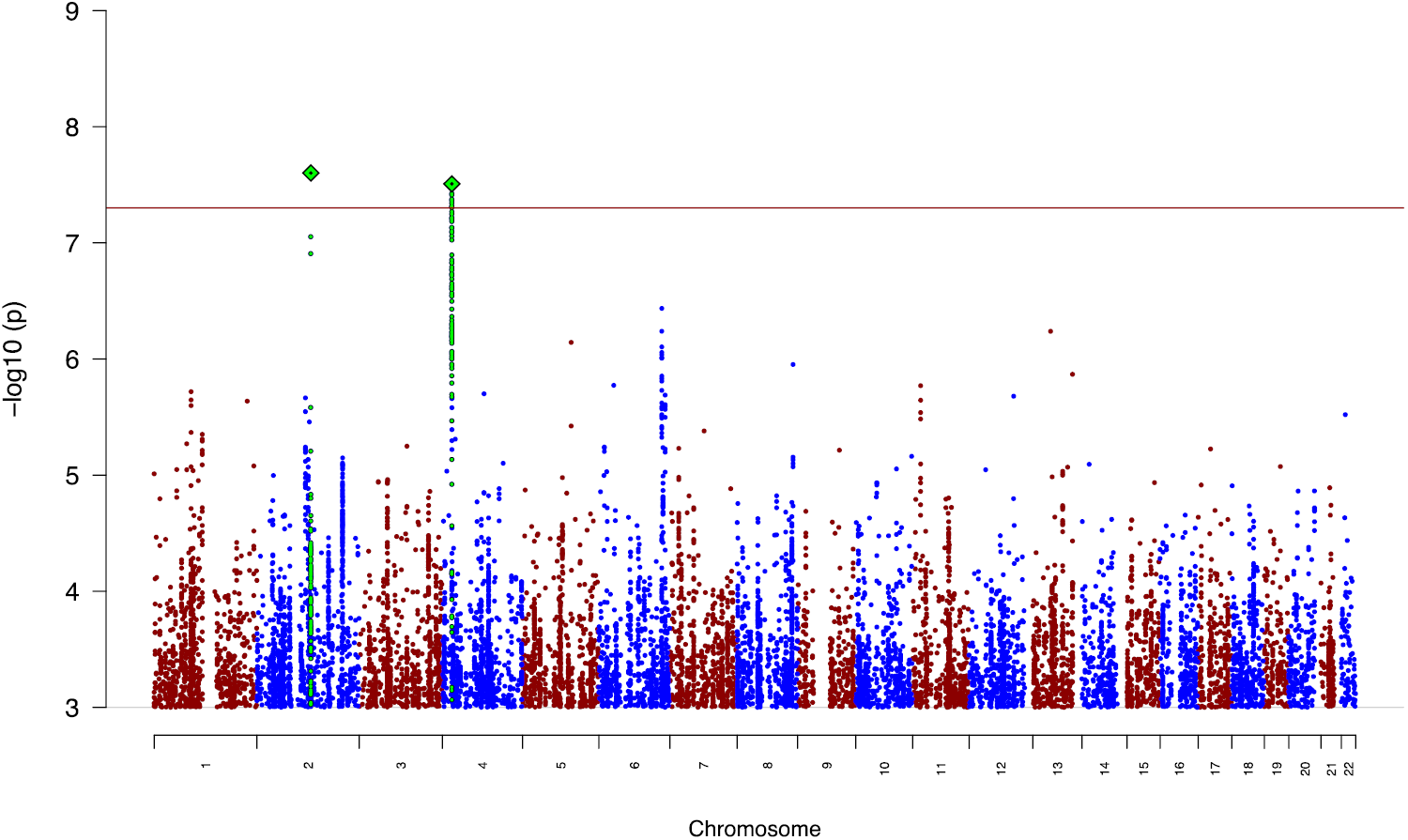
Manhattan plot for meta-analysis of suicide attempt in mood disorders. The red line shows the genome-wide significance threshold (P < 5 x 10^-8^). SNPs in green are in linkage disequilibrium with the index SNPs (diamonds).

### Replication

SNPs from the three genome-wide significant loci for suicide attempt in the discovery phase were tested for replication in independent mood disorder cohorts from the UK Biobank and *i*PSYCH. None of these loci showed association with suicide attempt in mood disorders in either study (minimum P value 0.22; Supplementary Table 14).

### Polygenic risk scoring and SNP-heritability

Polygenic risk scoring was performed to investigate the genetic etiology of suicide attempt in MDD, BIP and SCZ using scores from GWAS on psychiatric disorders and the SA GWAS conducted here. PRS for major depression were significantly associated with suicide attempt in all three disorders (MDD - maximum variance explained (R^2^) = 0.18%, P = 0.0006; BIP - R^2^ = 0.20%, P = 0.0002; SCZ - R^2^ = 0.33%, P = 0.0006; Figure 2). The PRS for BIP did not differ between suicide attempters and non-attempters with BIP, while PRS for schizophrenia were significantly lower in suicide attempters with schizophrenia compared with non-attempters (R^2^ = 0.35%, P = 0.0004, Supplementary Figure 7).

**Figure 2:**
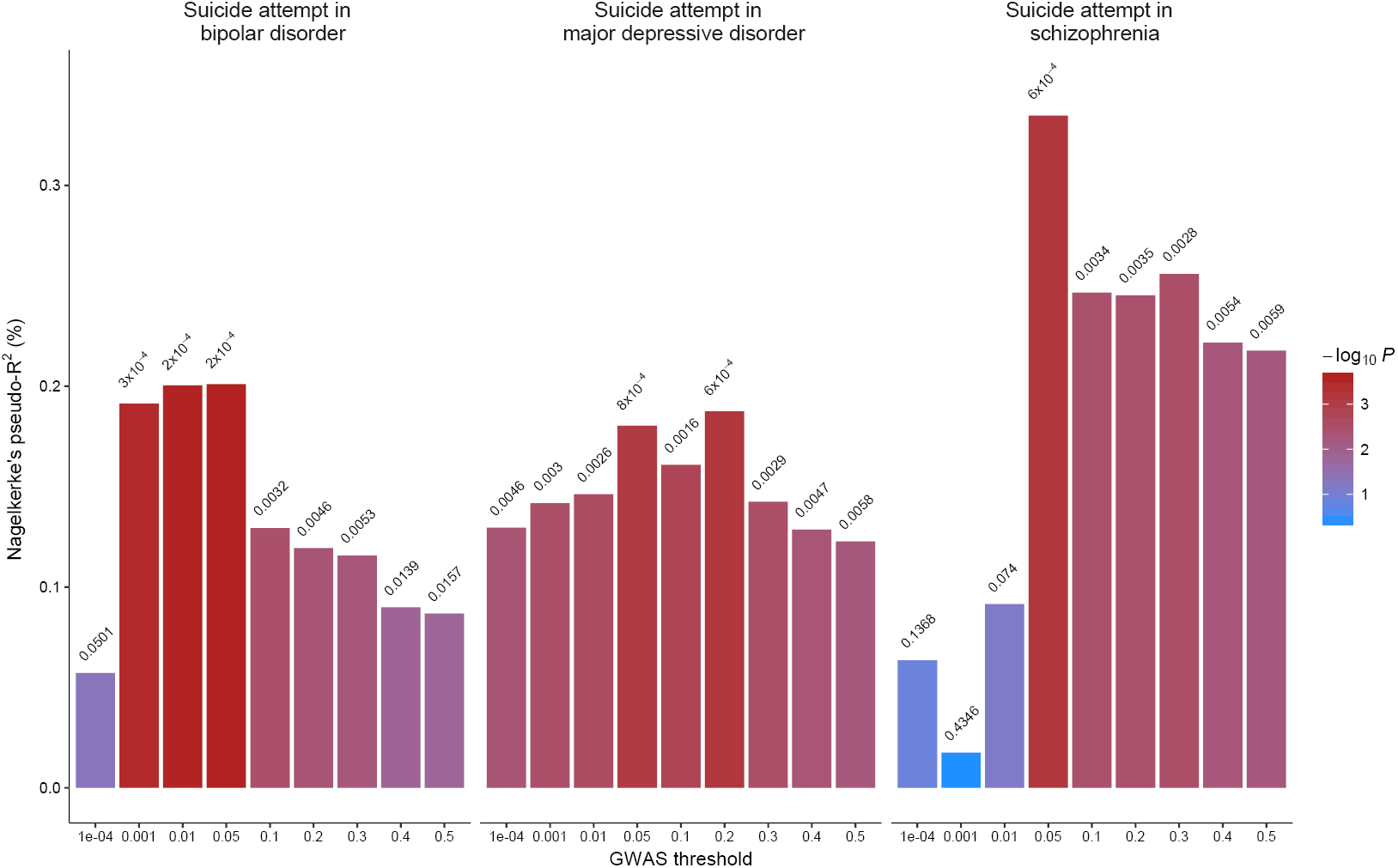
Polygenic risk scores for major depression are associated with suicide attempt versus non-attempt in bipolar disorder, major depressive disorder and schizophrenia. The x-axis shows the P value threshold used to select SNPs from the discovery GWAS. The y-axis shows the Nagelkerke’s pseudo-R^2^ measure of variance explained. P values of association between polygenic scores and suicide attempt are shown above each bar.

The SNP heritability 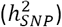 of suicide attempt in each psychiatric disorder was estimated using GREML. The 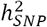estimate of suicide attempt in MDD was 0.03 (SE = 0.03, P = 0.19), in BIP was 0.02 (SE = 0.03, P = 0.25) and in SCZ was 0.10 (SE = 0.07, P = 0.06). None of these 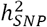estimates were significantly different from zero. Using these SA GWAS as discovery studies, the polygenic risk scores for SA in one psychiatric disorder were not associated with suicide attempt in another disorder (Supplementary Figure 6).

## Discussion

This GWAS on suicide attempt is the largest performed to date, combining samples of suicide attempters and non-attempters across three major psychiatric disorders and 46 individual cohorts from the Psychiatric Genomics Consortium. Three independent loci were associated with suicide attempt in the discovery phase. The strongest support was for the chromosome 4 locus in *LOC105374524*, a non-coding RNA, located approximately 500Kb downstream of the *GBA3* (Glucosylceramidase Beta 3) gene and upstream of *PPARGC1A*, which is a transcriptional coactivator that regulates the genes involved in energy metabolism. This region reached genome-wide significance in the GWAS of suicide attempt in bipolar disorder and the association strengthened in the meta-analysis of suicide attempt in mood disorders. Despite the support for this locus in the discovery phase, the association was not replicated in large independent mood disorder cohorts from the UK Biobank and *i*PSYCH. One possible explanation for this lack of replication is that over 90% of suicide attempters in the replication cohorts had a depression diagnosis, while in the discovery PGC studies, this locus had strongest effect in BIP. In addition, there may be heterogeneity in the definition of suicide attempt, which in the discovery samples was based on psychiatric interviews and in the replication samples was based on self-report questionnaires and hospital records. For the other two loci reaching genome-wide significance for suicide attempt in the discovery phase, statistical power for replication was low, given the effect allele frequency of 2% at each locus.

The within-case analysis strategy utilised in this GWAS was designed to detect associations specific for suicide attempt and was informed by twin and family studies which consistently indicate a genetic component of SA which is distinct from that of the psychiatric disorders themselves (5, 6). The *LOC105374524* association with suicide attempt in mood disorders reported here has not been implicated in major depression, BIP or SCZ in the latest GWAS conducted by the PGC, providing support for this study design (17-19).

The polygenic risk scoring analyses showed that suicide attempters with a MDD, BIP or SCZ diagnosis carry a greater liability for major depression than non-attempters. These results indicate the existence of a shared genetic etiology between SA and major depression that is common to suicide attempt in different psychiatric disorders. Clincal studies provide support for these findings, with the presence of depressive symptoms in schizophrenia and increased depressive symptoms in bipolar disorder being risk factors for suicide attempt (35, 36).

The SNP-heritability estimates for suicide attempt in each disorder were not significantly different from zero and while several twin studies have reported that suicide attempt is moderately heritable, one study found that after adjusting for psychiatric disorders, the heritability decreased from 30% to 17% (5, 37). Given estimated SNP-heritabilities are generally much lower than twin heritabilities, our samples were underpowered to identify this level of genetic contribution. If this is a more accurate estimate of the independent genetic contribution to suicide attempt, then substantial increases in sample size will be required to fully interrogate the genetic etiology of suicide attempt. Still, the present study is the first consortium-based GWAS on suicide attempt and makes significant progress in increasing numbers by combining samples across clinical cohorts. Further collaborative efforts to amass samples on an even larger scale will be essential to achieve the success seen in GWAS of other psychiatric disorders. Data from population biobanks are now widely available and leveraging these to conduct meta-analyses could rapidly increase statistical power. GWAS comparing suicide attempters versus healthy controls may provide a complementary approach by prioritising loci for follow-up in case-only studies, which should be expanded beyond studies of mood disorders and schizophrenia to also include existing cohorts for other disorders where suicide attempt is prevalent. The results of GWAS now robustly link hundreds of genetic loci to psychiatric disorders and provide additional opportunities to disentangle genetic effects on suicide attempt from the disorders themselves.

One strength of this study is that samples of suicide attempters have been successfully combined across many individual clinical cohorts. All 46 datasets were processed centrally using the same quality control, imputation and analysis pipeline. Suicide attempt and non-attempt were defined using items from structured psychiatric interviews, although it should be noted that these items vary by interview which could result in heterogeneity in the phenotype definition. Since subjects were not ascertained primarily for suicide attempt, detailed information such as the number of suicide attempts, medical consequences or medication is not available for all participants. This study focused on lifetime suicide attempt to maximize sample size, but some cases who were non-attempters at the time of recruitment may later have attempted suicide.

Of all the psychiatric phenotypes, suicidality remains especially challenging to predict and assess and there is an urgent need to better understand its etiology. As seen in GWAS of other psychiatric disorders, the number of genetic associations is expected to accumulate with increased sample size and can provide invaluable biological insights. Our novel finding that genetic liability for major depression increases risk of suicide attempt in MDD, BIP, and SCZ provides a possible starting point for predictive modelling and preventative strategies. The ultimate goal of genetic studies on suicide attempt is to translate these statistical associations into biological mechanisms and much-needed treatments and preventions for suicidality, in order to reduce its burden on patients, families and healthcare systems.

## Acknowledgements

We are deeply indebted to the investigators who comprise the PGC, and to the subjects who have shared their life experiences with PGC investigators. The PGC has received major funding from the US National Institute of Mental Health and the US National Institute of Drug Abuse (U01 MH109528 and U01 MH1095320). Statistical analyses were carried out on the Genetic Cluster Computer (http://www.geneticcluster.org) hosted by SURFsara and financially supported by the Netherlands Scientific Organization (NWO 480-05-003 PI: Posthuma) along with a supplement from the Dutch Brain Foundation and the VU University Amsterdam. This research has been conducted using the UK Biobank Resource (http://www.ukbiobank.ac.uk/), as an approved extension to application 16577 (Dr Breen). The iPSYCH team acknowledges funding from the Lundbeck Foundation (grants R102-A9118 and R155-2014-1724), the Stanley Medical Research Institute, the European Research Council (project 294838), the Novo Nordisk Foundation for supporting the Danish National Biobank resource, and Aarhus and Copenhagen Universities and University Hospitals, including support to the iSEQ Center, the GenomeDK HPC facility, and the CIRRAU Center. Some data used in this study were obtained from dbGaP. Funding support for the Genome-Wide Association of Schizophrenia Study was provided by the National Institute of Mental Health (R01 MH67257, R01 MH59588, R01 MH59571, R01 MH59565, R01 MH59587, R01 MH60870, R01 MH59566, R01 MH59586, R01 MH61675, R01 MH60879, R01 MH81800, U01 MH46276, U01 MH46289 U01 MH46318, U01 MH79469, and U01 MH79470) and the genotyping of samples was provided through the Genetic Association Information Network (GAIN). The datasets used for the analyses described in this manuscript were obtained from the database of Genotypes and Phenotypes (dbGaP) found at http://www.ncbi.nlm.nih.gov/gap through dbGaP accession number phs000021.v3.p2. Samples and associated phenotype data for the Genome-Wide Association of Schizophrenia Study were provided by the Molecular Genetics of Schizophrenia Collaboration (PI: Pablo V. Gejman, Evanston Northwestern Healthcare (ENH) and Northwestern University, Evanston, IL, USA).” Funding support for the Whole Genome Association Study of Bipolar Disorder was provided by the National Institute of Mental Health (NIMH) and the genotyping of samples was provided through the Genetic Association Information Network (GAIN). The datasets used for the analyses described in this manuscript were obtained from the database of Genotypes and Phenotypes (dbGaP) found at http://www.ncbi.nlm.nih.gov/gap through dbGaP accession number phs000017.v3.p1. Samples and associated phenotype data for the Collaborative Genomic Study of Bipolar Disorder were provided by the The NIMH Genetics Initiative for Bipolar Disorder. Data and biomaterials were collected in four projects that participated in NIMH Bipolar Disorder Genetics Initiative. From 1991-98, the Principal Investigators and Co-Investigators were: Indiana University, Indianapolis, IN, U01 MH46282, John Nurnberger, M.D., Ph.D., Marvin Miller, M.D., and Elizabeth Bowman, M.D.; Washington University, St. Louis, MO, U01 MH46280, Theodore Reich, M.D., Allison Goate, Ph.D., and John Rice, Ph.D.; Johns Hopkins University, Baltimore, MD U01 MH46274, J. Raymond DePaulo, Jr., M.D., Sylvia Simpson, M.D., MPH, and Colin Stine, Ph.D.; NIMH Intramural Research Program, Clinical Neurogenetics Branch, Bethesda, MD, Elliot Gershon, M.D., Diane Kazuba, B.A., and Elizabeth Maxwell, M.S.W. Data and biomaterials were collected as part of ten projects that participated in the NIMH Bipolar Disorder Genetics Initiative. From 1999-03, the Principal Investigators and Co-Investigators were: Indiana University, Indianapolis, IN, R01 MH59545, John Nurnberger, M.D., Ph.D., Marvin J. Miller, M.D., Elizabeth S. Bowman, M.D., N. Leela Rau, M.D., P. Ryan Moe, M.D., Nalini Samavedy, M.D., Rif El-Mallakh, M.D. (at University of Louisville), Husseini Manji, M.D. (at Wayne State University), Debra A. Glitz, M.D. (at Wayne State University), Eric T. Meyer, M.S., Carrie Smiley, R.N., Tatiana Foroud, Ph.D., Leah Flury, M.S., Danielle M. Dick, Ph.D., Howard Edenberg, Ph.D.; Washington University, St. Louis, MO, R01 MH059534, John Rice, Ph.D, Theodore Reich, M.D., Allison Goate, Ph.D., Laura Bierut, M.D.; Johns Hopkins University, Baltimore, MD, R01 MH59533, Melvin McInnis M.D., J. Raymond DePaulo, Jr., M.D., Dean F. MacKinnon, M.D., Francis M. Mondimore, M.D., James B. Potash, M.D., Peter P. Zandi, Ph.D, Dimitrios Avramopoulos, and Jennifer Payne; University of Pennsylvania, PA, R01 MH59553, Wade Berrettini M.D.,Ph.D.; University of California at Irvine, CA, R01 MH60068, William Byerley M.D., and Mark Vawter M.D.; University of Iowa, IA, R01 MH059548, William Coryell M.D., and Raymond Crowe M.D.; University of Chicago, IL, R01 MH59535, Elliot Gershon, M.D., Judith Badner Ph.D., Francis McMahon M.D., Chunyu Liu Ph.D., Alan Sanders M.D., Maria Caserta, Steven Dinwiddie M.D., Tu Nguyen, Donna Harakal; University of California at San Diego, CA, R01 MH59567, John Kelsoe, M.D., Rebecca McKinney, B.A.; Rush University, IL, R01 MH059556, William Scheftner M.D., Howard M. Kravitz, D.O., M.P.H., Diana Marta, B.S., Annette Vaughn-Brown, MSN, RN, and Laurie Bederow, MA; NIMH Intramural Research Program, Bethesda, MD, 1Z01MH002810-01, Francis J. McMahon, M.D., Layla Kassem, PsyD, Sevilla Detera-Wadleigh, Ph.D, Lisa Austin,Ph.D, Dennis L. Murphy, M.D.

## References

1. World Health Organization. Preventing suicide: A global imperative. Geneva 2014.

2. Centers for Disease Control and Prevention. Suicide: Facts at a Glance 2015 [Available from: https://www.cdc.gov/violenceprevention/pdf/suicide-datasheet-a.pdf.

3. Qin P. The impact of psychiatric illness on suicide: differences by diagnosis of disorders and by sex and age of subjects. J Psychiatr Res. 2011;45(11):1445–52.

4. Beautrais AL, Joyce PR, Mulder RT, Fergusson DM, Deavoll BJ, Nightingale SK. Prevalence and comorbidity of mental disorders in persons making serious suicide attempts: a case-control study. Am J Psychiatry. 1996;153(8):1009–14.

5. Voracek M, Loibl LM. Genetics of suicide: a systematic review of twin studies. Wien Klin Wochenschr. 2007;119(15-16):463–75.

6. Brent DA, Mann JJ. Family genetic studies, suicide, and suicidal behavior. Am J Med Genet C Semin Med Genet. 2005;133C(1):13–24.

7. Willour VL, Seifuddin F, Mahon PB, Jancic D, Pirooznia M, Steele J, et al. A genome-wide association study of attempted suicide. Mol Psychiatry. 2012;17(4):433–44.

8. Schosser A, Butler AW, Ising M, Perroud N, Uher R, Ng MY, et al. Genomewide association scan of suicidal thoughts and behaviour in major depression. Plos One. 2011;6(7):e20690.

9. Perlis RH, Huang J, Purcell S, Fava M, Rush AJ, Sullivan PF, et al. Genome-wide association study of suicide attempts in mood disorder patients. Am J Psychiatry. 2010;167(12):1499–507.

10. Mullins N, Perroud N, Uher R, Butler AW, Cohen-Woods S, Rivera M, et al. Genetic relationships between suicide attempts, suicidal ideation and major psychiatric disorders: a genome-wide association and polygenic scoring study. Am J Med Genet B Neuropsychiatr Genet. 2014;165B(5):428–37.

11. Galfalvy H, Haghighi F, Hodgkinson C, Goldman D, Oquendo MA, Burke A, et al. A genome-wide association study of suicidal behavior. Am J Med Genet B Neuropsychiatr Genet. 2015;168(7):557–63.

12. Stein MB, Ware EB, Mitchell C, Chen CY, Borja S, Cai T, et al. Genomewide association studies of suicide attempts in US soldiers. Am J Med Genet B Neuropsychiatr Genet. 2017;174(8):786–97.

13. Ruderfer DM, Walsh CG, Aguirre MW, Ribeiro JD, Franklin JC, Rivas MA. Significant shared heritability underlies suicide attempt and clinically predicted probability of attempting suicide. bioRxiv. 2018.

14. American Psychiatric Association. Diagnostic and Statistical Manual of Mental Disorders 4th edition Washington, DC: American Psychiatric Association,; 1994.

15. World Health Organization. International Classification of Diseases 9th revised edn. 9th revised edn ed: World Health Organization; 1978.

16. World Health Organization. International Classification of Diseases 10th revised edn. 10th revised edn ed: World Health Organization; 1992.

17. Wray NR, Ripke S, Mattheisen M, Trzaskowski M, Byrne EM, Abdellaoui A, et al. Genome-wide association analyses identify 44 risk variants and refine the genetic architecture of major depression. Nat Genet. 2018;50(5):668–81.

18. Stahl E, Forstner A, McQuillin A, Ripke S, Bipolar Disorder Working Group of the PGC, Ophoff R, et al. Genomewide association study identifies 30 loci associated with bipolar disorder. bioRxiv. 2017.

19. Schizophrenia Working Group of the Psychiatric Genomics Consortium. Biological insights from 108 schizophrenia-associated genetic loci. Nature. 2014;511(7510):421–7.

20. Howie B, Marchini J, Stephens M. Genotype imputation with thousands of genomes. G3 (Bethesda). 2011;1(6):457–70.

21. Delaneau O, Marchini J, Zagury JF. A linear complexity phasing method for thousands of genomes. Nat Methods. 2011;9(2):179–81.

22. 1000 Genomes Project Consortium. A map of human genome variation from population-scale sequencing. Nature. 2010;467(7319):1061–73.

23. Chang CC, Chow CC, Tellier LC, Vattikuti S, Purcell SM, Lee JJ. Second-generation PLINK: rising to the challenge of larger and richer datasets. GigaScience. 2015;4:7.

24. Price AL, Patterson NJ, Plenge RM, Weinblatt ME, Shadick NA, Reich D. Principal components analysis corrects for stratification in genome-wide association studies. Nat Genet. 2006;38(8):904–9.

25. Willer CJ, Li Y, Abecasis GR. METAL: fast and efficient meta-analysis of genomewide association scans. Bioinformatics. 2010;26(17):2190–1.

26. Euesden J, Lewis CM, O’Reilly PF. PRSice: Polygenic Risk Score software. Bioinformatics. 2015;31(9):1466–8.

27. Yang J, Lee SH, Goddard ME, Visscher PM. GCTA: a tool for genome-wide complex trait analysis. Am J Hum Genet. 2011;88(1):76–82.

28. Purcell S, Cherny SS, Sham PC. Genetic Power Calculator: design of linkage and association genetic mapping studies of complex traits. Bioinformatics. 2003;19(1):149–50.

29. Visscher PM, Hemani G, Vinkhuyzen AA, Chen GB, Lee SH, Wray NR, et al. Statistical power to detect genetic (co)variance of complex traits using SNP data in unrelated samples. Plos Genet. 2014;10(4):e1004269.

30. Dudbridge F. Power and predictive accuracy of polygenic risk scores. Plos Genet. 2013;9(3):e1003348.

31. Palla L, Dudbridge F. A Fast Method that Uses Polygenic Scores to Estimate the Variance Explained by Genome-wide Marker Panels and the Proportion of Variants Affecting a Trait. Am J Hum Genet. 2015;97(2):250–9.

32. Sudlow C, Gallacher J, Allen N, Beral V, Burton P, Danesh J, et al. UK biobank: an open access resource for identifying the causes of a wide range of complex diseases of middle and old age. Plos Med. 2015;12(3):e1001779.

33. Bycroft C, Freeman C, Petkova D, Band G, Elliott LT, Sharp K, et al. Genome-wide genetic data on ∼500,000 UK Biobank participants. bioRxiv. 2017.

34. Pedersen CB, Bybjerg-Grauholm J, Pedersen MG, Grove J, Agerbo E, Baekvad-Hansen M, et al. The iPSYCH2012 case-cohort sample: new directions for unravelling genetic and environmental architectures of severe mental disorders. Mol Psychiatry. 2018;23(1):6–14.

35. Marangell LB, Bauer MS, Dennehy EB, Wisniewski SR, Allen MH, Miklowitz DJ, et al. Prospective predictors of suicide and suicide attempts in 1,556 patients with bipolar disorders followed for up to 2 years. Bipolar Disord. 2006;8(5 Pt 2):566–75.

36. Hawton K, Sutton L, Haw C, Sinclair J, Deeks JJ. Schizophrenia and suicide: systematic review of risk factors. Br J Psychiatry. 2005;187:9–20.

37. Fu Q, Heath AC, Bucholz KK, Nelson EC, Glowinski AL, Goldberg J, et al. A twin study of genetic and environmental influences on suicidality in men. Psychol Med. 2002;32(1):11– 24.

